# Yeast nicotinate-nucleotide pyrophosphorylase in complex with its ligand: Crystallization and Preliminary structural approaches

**DOI:** 10.1101/2022.11.23.517371

**Authors:** Tae Gyun Kim, Taek Hun Kwon, Hyo Jung Choi, Min Kyung Park, Jungbae Park, Inkyung Chae, Hyun-Jung An

## Abstract

Pyridine-2,3-dicarboxylic acid which is a biologically potent molecule implicated in neurodegenerative environment is catalyzed by nicotinate-nucleotide pyrophosphorylase (NMnPP) to produce a precursor molecule, nicotinate mononucleotide (NMn), of *de novo* biosynthesis of the coenzyme nicotinamide adenine dinucleotide (NAD^+^). The protein preparation, crystallization, and preliminary structural features of full-length enzyme in complex with product reactant suggest that yeast NMnPP acts as stable hexamer formation. *S. cerevisiae* NMnPP was obtained and diffracted to a resolution of 1.74 Å and 1.99 Å for apo and complex forms, belonged to the trigonal symmetry group *R*32 in the unit-cell parameters of a=b=155.313, c=67.507 and a=b=155.091, c=69.204, respectively. Based on our comparison of eukaryotic NMnPP structures in the apo and complex forms, we propose functional and structural investigation for the product binding and hexamer stabilization.

## INTRODUCTION

NAD^+^ is crucial to cellular energy metabolism, the fermentation cycle, and oxidative phosphorylation in the respiratory system (Amjad *et al*., 2021). Pyridine-2,3-dicarboxylic acid, also known as quinolinic acid (QA), is catabolized by NMnPP in the first step of *de novo* NAD^+^ metabolite and kynurenine pathway (Fig. 1) and is a potent excitotoxic compound in the central nervous system that may be involved in psychiatric disorders, the neurodegenerative process in the brain, and acts as an agonist of N-methyl-D-aspartate receptors (Vega-Naredo *et al*., 2005). NMnPP plays an important role as QA homeostasis in the brain, liver, and kidney (Braidy *et al*., 2011). Malfunction of NMnPP results from an accumulation of QA levels, which causes strongly involved in a series of severe neurodegenerative disorders including Huntington’s disease, Alzheimer’s disease, and AIDS dementia complex (Beal *et al*., 1991; Guillemin *et al*., 2005a & 2005b). NMnPP as a potential therapeutic target in malignant gliomas, which is exclusively expressed in the WHO high-grade III–IV glioma, exploits microglia-derived QA as an alternative source of replenishing metabolic NAD+. This process enables glioma to survive under conditions of oxidative stress and NAD+ loss caused by therapeutic approaches, including alkylating agents and direct NAD+ synthesis inhibitors (Opitz *et al*., 2011; Sahm *et al*., 2013).

**Figure 1.**
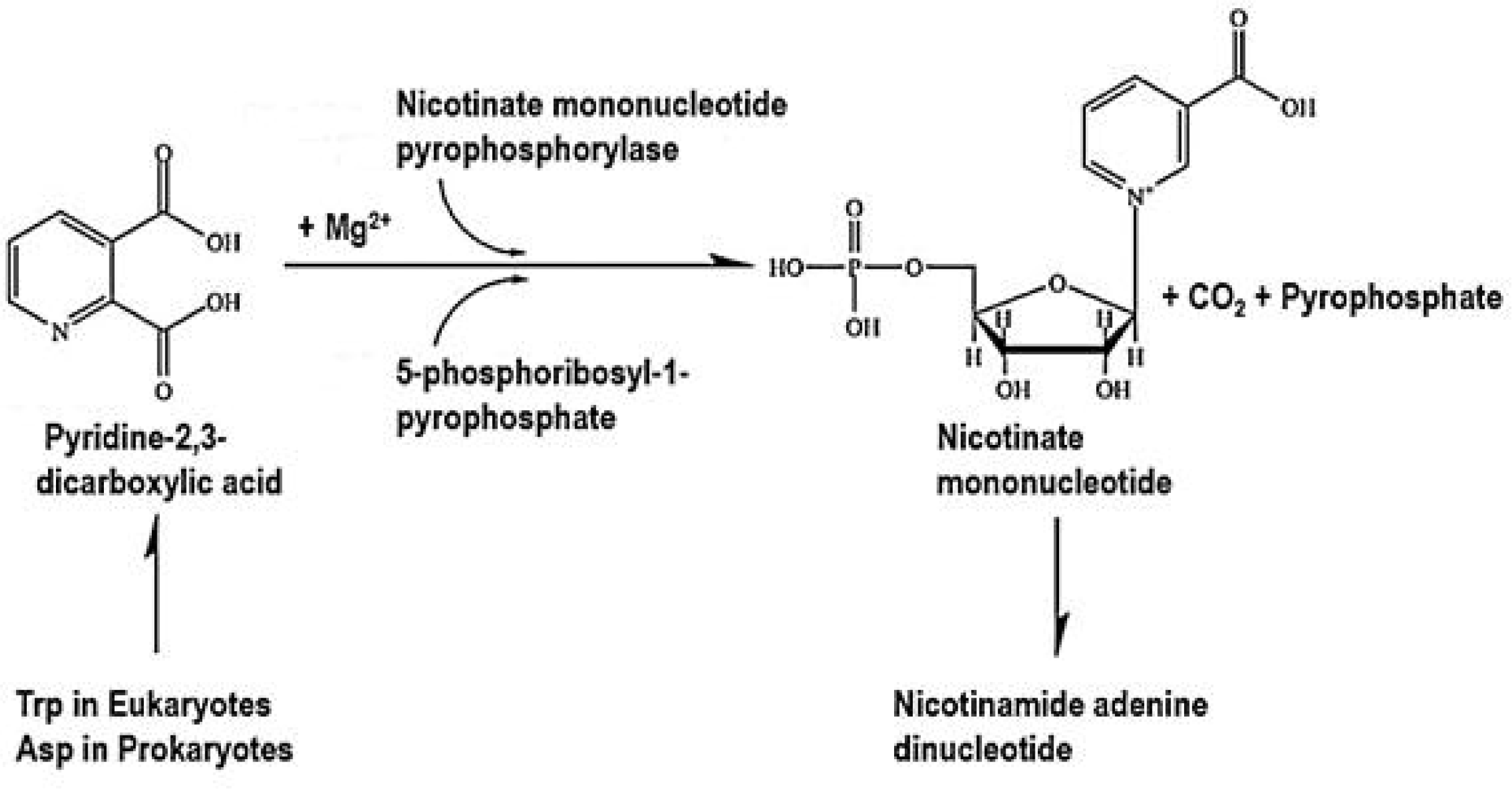
The catalytic reaction of nicotinate mononucleotide pyrophosphorylase (NMnPP).

NMnPP structures from eukaryotes to prokaryotes have been reported as dimers or hexamers. Based on structural and functional comparisons of key enzymes in the kynurenine pathway, the amino acid sequence of *Saccharomyces cerevisiae* (Uniprot # P43619) shows the average sequence identity of 40.8% to that of well-known NMmPP from *Homo sapiens* (Uniprot # Q15274), *Sus scrofa* (Uniprot # I3LK75), *Bos taurus* (Uniprot # Q3T063), *Helicobacter pylori* (Uniprot # Q9ZJN2), *Mycobacterium tuberculosis* (Uniprot # P9WJJ7), *Salmonella typhimurium* (Uniprot # P30012), *Pseudomonas aeruginosa* (Uniprot # P30819), and *Escherichia coli* (Uniprot # P30011) (Fig. 2). All structures exhibit typical features of the type II pyrophosphorylase structure including an N-terminal four-stranded β-sandwich domain and a C-terminal α/β barrel domain, unlike the α4/β5 folds of other types of pyrophosphorylase (Youn *et al*., 2013,2016; Kim *et al*., 2006,2007; Liu *et al*., 2007; Di Luccio *et al*., 2008). Whereas most prokaryotic NMnPP has a dimeric formation, eukaryotic NMnPP included some NMnPP in prokaryotes such as *H. pylori* and *Thermus thermophilus* (reported in PDB 1×1o) exists as hexamers. The dimer-dimer interfaces by *H. pylori* NMnPP mutant as F181P and N-terminal truncated form of human NMnPP showed the oligomeric status from hexamer to dimer, which result from enzyme stability and non-functionality (Youn *et al*., 2016; Kim *et al*., 2007). Although numerous NMnPP structures have been determined in apo and complex with ligands, the structural and functional basis on the hexamer moieties of NMnPP still remain unclear.

**Figure 2.**
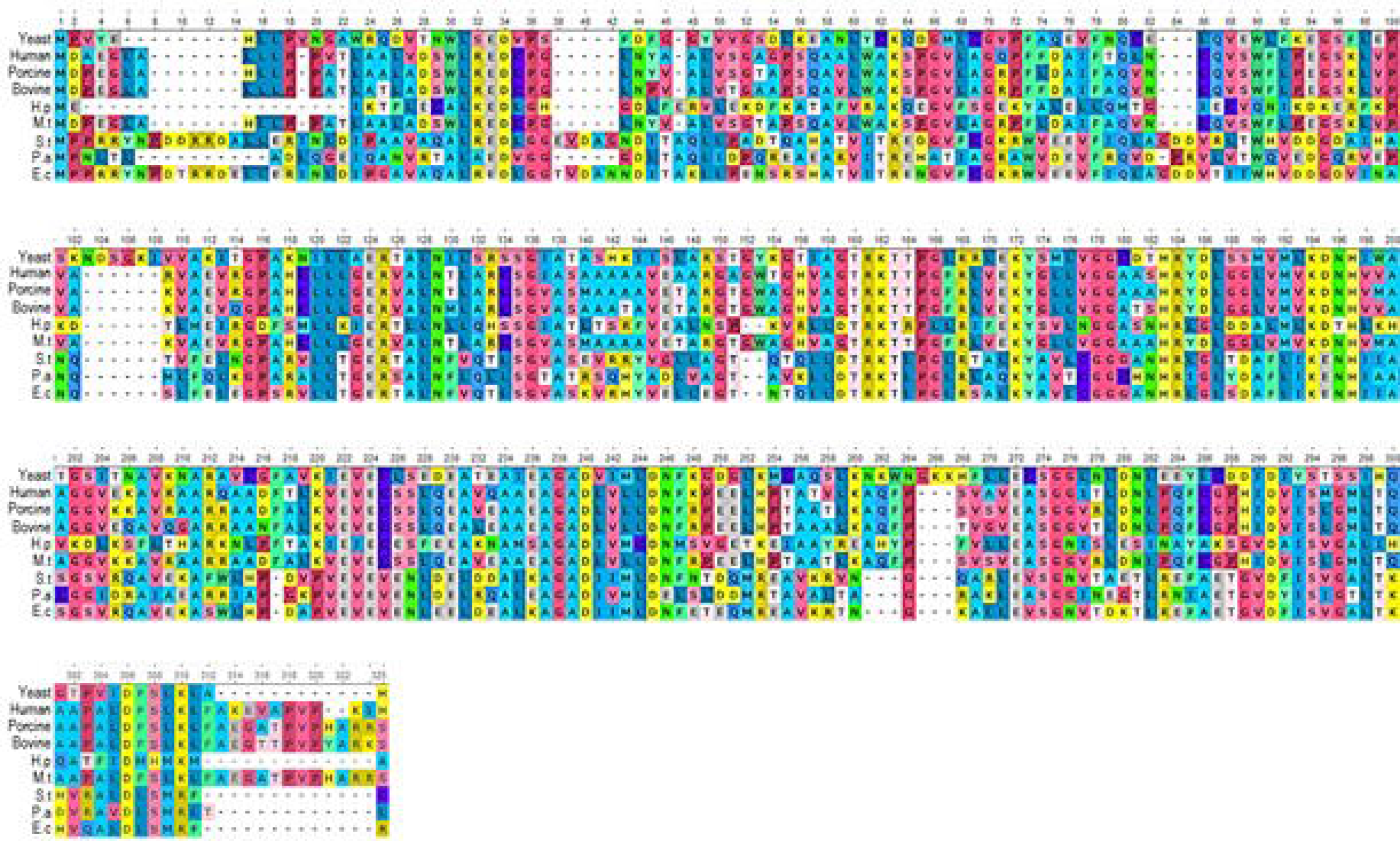
The amino acid sequence alignment among yeast NMnPP and other NMnPPs was generated and modified by multiple alignment program Unipro UGENE. The accession number and database of aligned sequences are indicated in parentheses: Yeast, *Saccharomyces cerevisiae* (Uniprot # P43619); Human, *Homo sapiens* (Uniprot # Q15274); Porcine, *Sus scrofa* (Uniprot # I3LK75); Bovine, *Bos taurus* (Uniprot # Q3T063); H.p, *Helicobacter pylori* (Uniprot # Q9ZJN2); M.t, *Mycobacterium tuberculosis* (Uniprot # P9WJJ7); S.t, *Salmonella typhimurium* (Uniprot # P30012); P.a, *Pseudomonas aeruginosa* (Uniprot # P30819); E.c, *Escherichia coli* (Uniprot # P30011).

Here, we preliminarily analysed the structural solution on apo and NMm-bound forms of NMmPP at high resolution using X-ray crystallographic and electron microscopic techniques. Reproducible crystals for apo and NMm-bound NMmPP were obtained and diffracted to 1.74 and 1.99 Å resolution, respectively. Our results provide structural insight into the crucial information of the product binding and hexamer stabilisation of eukaryotic NMmPP and suggest clinical implications in the development of anticancer agents on the therapeutic direction of severe neurodegenerative malfunction.

## RESULTS AND DISCUSSION

The recombinant nicotinate-nucleotide pyrophosphorylase (NMnPP) protein with an N-terminal TEV protease-cleavable 6xHis-tag was successfully overexpressed as a soluble form in *E. coli* C43(DE3) and purified to crystallizable quality using Ni-NTA affinity chromatography, followed by cleavage of the N-terminal His-tag with TEV protease, MonoQ ion exchange chromatography, and HiLoad Superdex 200 20/60 prep grade chromatography (GE Healthcare, USA). The final yield of NMnPP protein was concentrated at approximately 15 mg per L of cultured cells, and the highly qualified purity of the protein sample was judged by Coomassie-stained SDS-PAGE, indicating as a monomeric 21 kDa on SDS-PAGE used a protein marker (Bio-rad) (Fig. 3A).

**Figure 3.**
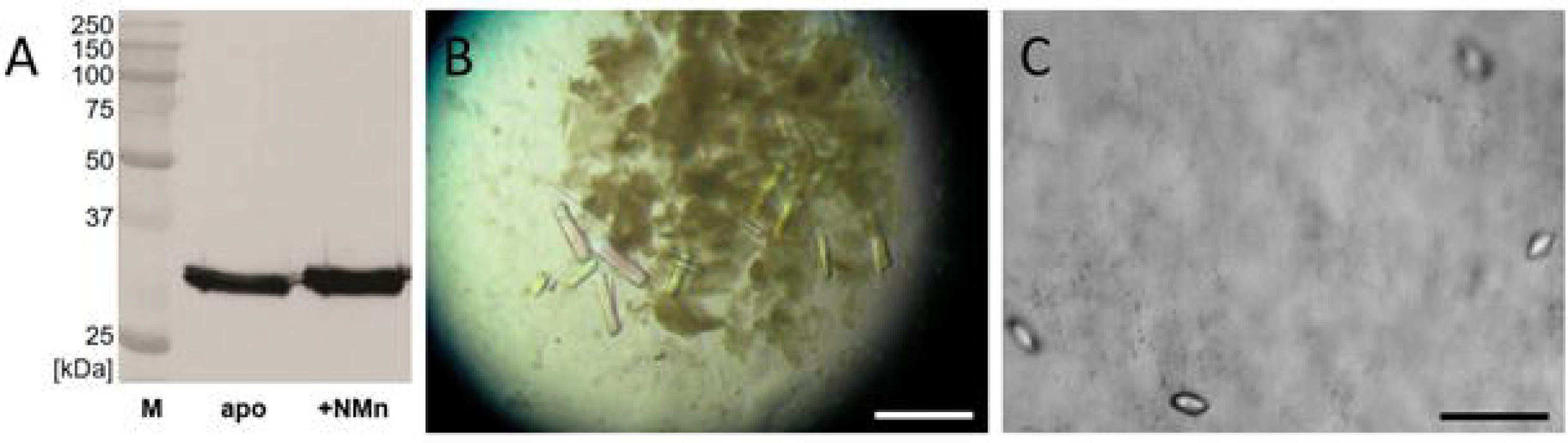
Purified NMnPP on a 18% SDS-PAGE (A). Lane M, molecular-weight markers; apo, final purified NMnPP; +NMn, a mixture of NMn with NMnPP. The apo and NMn-bound NMnPP crystals were obtained as rod-like (B) and rice-like (C) forms, respectively. The scale bar indicates 200 μm.

Initial crystallization of apo and NMn-bound NMnPP was carried out using the sitting-drop vapour diffusion method at 20°C. For diffraction-quality crystals, apo and NMn-bound NMnPP were obtained using 0.1 *M* Hepes-KOH, pH 7.3, 25% (w/v) PEG 3.35K, 0.2 *M* Mg(OAc)_2_ and 0.1 *M* Bis-tris propane, pH 6.5, 15% (v/v) MPD, 50 m*M* Na-Cacodylate, indicating that the concentration of MPD was critical to be crystallized as an optimized formation of NMn-bound NMnPP (Fig. 3B & 3C). After transferring apo and NMn-bound NMnPP crystals into 25% (v/v) glycerol in mother liquor for cryo-protection, optimized apo and NMn-bound NMnPP crystals were directly flash-cooled in a gas stream from liquid N_2_. Apo and NMn-bound crystals belonged to the trigonal symmetry space group *R*32 with four monomers per asymmetric unit. The Matthews coefficient and solvent content for apo and NMn-bound proteins were estimated to be 2.75 and 2.73 Å^3^ Da^−1^, corresponding to solvent contents of 55.34 and 55.01%, respectively (Matthews, 1968).

The molecular replacement was performed with MOLREP (Vagin and Teplyakov, 2010) and PHASER (McCoy *et al*., 2007) using the *S. cerevisiae* NMnPP structure (PDB code: 3C2E) as a search model. Initial refinement produced a possible model of apo NMnPP with R_work_ of 29.5% and R_free_ of 35.1% using PHENIX (Adams *et al*., 2010). The visualization of the electron density maps and manual build-up of the atomic models were carried out using the COOT program (v 9.6.0, Emsley and Cowtan, 2004). Further model building and refinement of the apo and NMn-bound structures was of well-qualified map to build a three-dimensional structure for apo and NMn-bound crystals diffracted to a resolution of 1.74 and 1.99 Å, respectively (Fig. 4). Statistics for diffraction data collection and processing details are summarized in Table 3. To investigate a molecular distribution pattern for NMmPP protein using negative-stain Transmission electron microscopic (TEM) method, we showed the structural morphology of apo and NMn-bound NMmPP proteins (Fig. 5A & 5B, respectively), indicating that conserved known NMmPP structures has a hexameric oligomerization.

**Figure 4.**
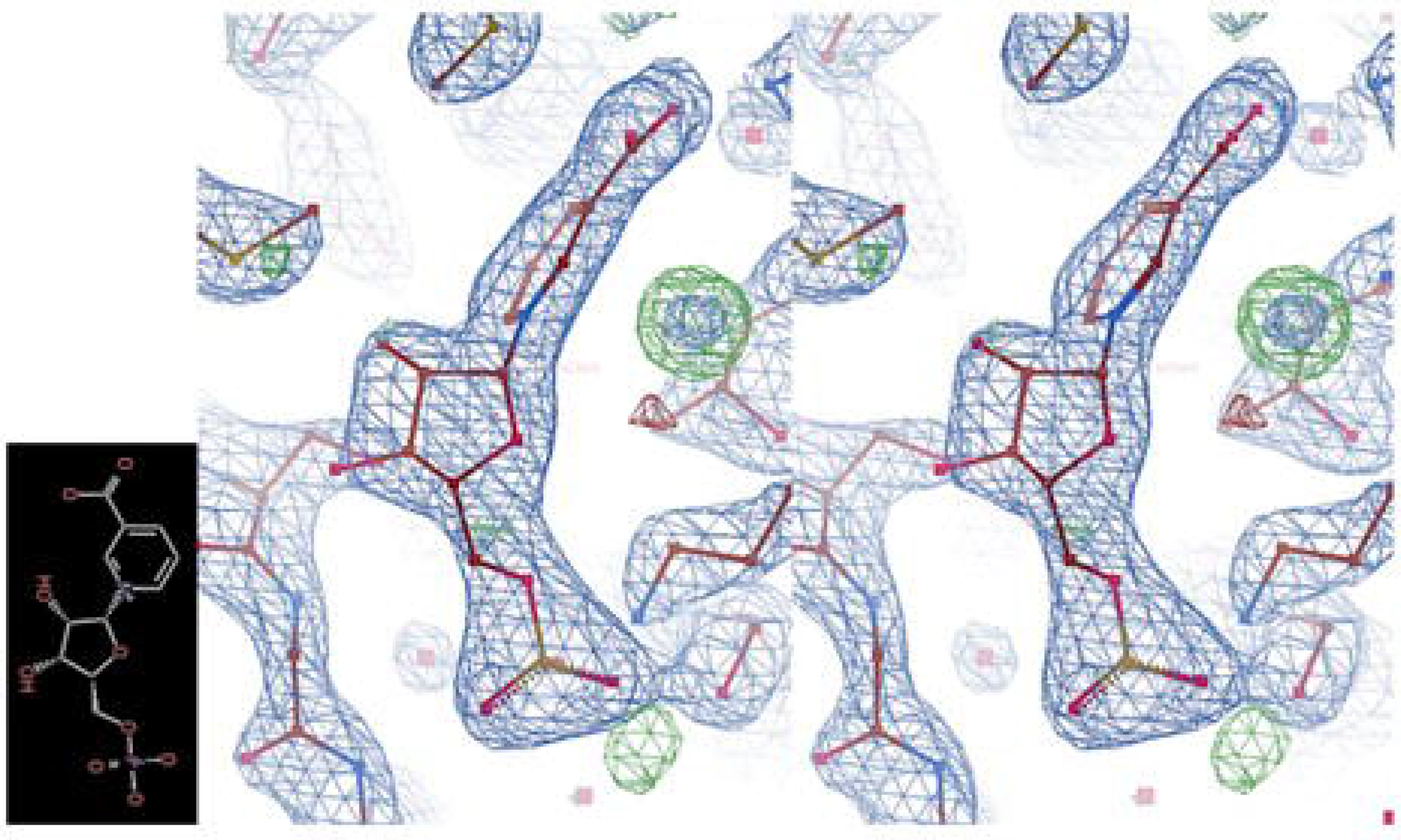
Initial 2Fo-Fc maps of NMn (chemical formula in left)-bound NMnPP structure. The structure is shown as a red linear stick and 2Fo-Fc maps (contoured at 1.49 σ) are stereo-viewed as blue meshes.

**Figure 5.**
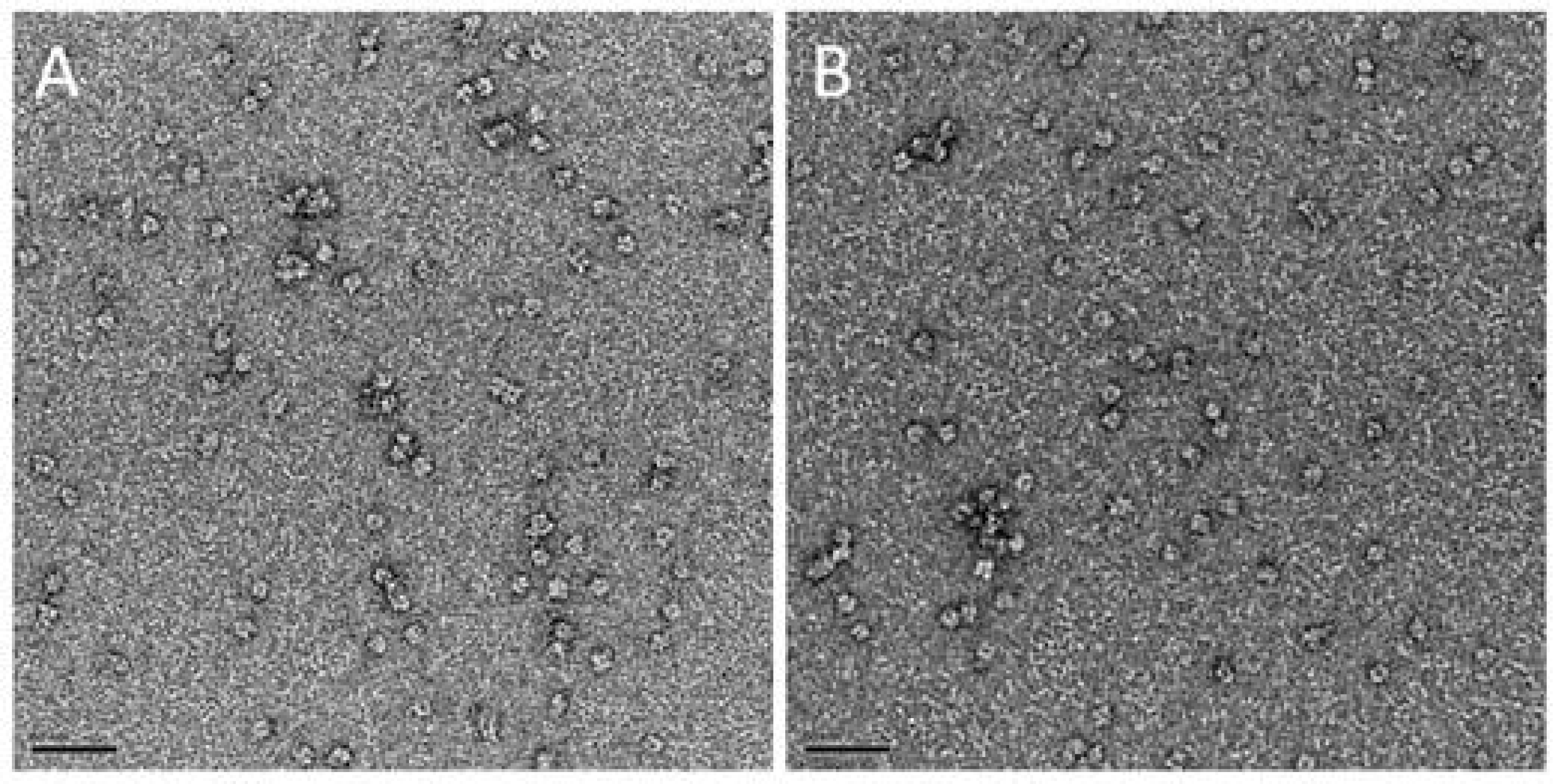
Negative-stain TEM analysis of apo (A) and NMn-bound NMnPP (B) proteins. The scale bar indicates 50 nm.

In this study, the purification, crystallization, and preliminary X-ray crystallographic and negative-stain TEM analyses provide a structural investigation of NMnPP to explain the further determination of flexible structural features compared to the current yeast NMnPP structure and the mechanical understanding of the kynurenine pathway of tryptophan degradation resulting from the *de novo* biosynthesis of NAD^+^.

## METHODS

### Protein expression and purification

Commercially available genomic DNA from *Saccharomyces cerevisiae* S288c was used as a template to amplify the nicotinate-nucleotide pyrophosphorylase (*Nmmpp*, Uniprot # P43619) gene by a polymerase chain reaction using indicated primers shown in Table 1. To confirm restriction enzyme sites that are excluded in the full coding sequence of *Nmmpp*, the uncleaved restriction enzyme site was checked with NEBcutter V2.0. We selected *Bam*HI and *Xho*I restriction sites of the expression vector pET-28a-TEV (Tobacco Etch Virus), which includes a cleavage site by TEV protease site in the N-terminus. To carry out to over-expressed cells, the pET-28a-TEV-*Nmmpp* was transformed and expressed in *E*. *coli* C43(DE3) (Invitrogen, USA) cells at 32°C with 0.4 m*M* Isopropyl-D-thiogalactoside (GE Healthcare) for 8 h. After collection of cultured cells by centrifugation, cell pellets were resuspended with buffer A (20m*M* Tris-HCl, pH 8.0, 300 m*M* NaCl, 5m*M* β-mercaptoethanol, 1mg/ml lysozyme, and mixture of protease inhibitors included in aprotinin, leupeptin, pepstatin, and PMSF) and gently disrupted by sonication (VC505, Sonics) on ice. Cell debris and supernatant were separated by high-speed centrifugation at 20,000 x g for 1 h at 4 °C, and the supernatant was collected. Impurities were removed by 0.45 μm filtration. The filtered supernatant included NMnPP protein was applied to a nickel-nitrilotriacetic acid (Ni-NTA, GE Healthcare) affinity column pre-equilibrated with buffer A. The Ni-NTA column was washed by using 50 times of the column volume by buffer A and 50m*M* Imidazole. Target protein was eluted with a linear gradient using buffer B (50 m*M* Tris-HCl, pH 8.0, 300 m*M* NaCl, 300 m*M* imidazole, and 5 m*M* β-mercaptoethanol). Eluted NMnPP protein was dialyzed against buffer A while simultaneously digesting with TEV protease for at least 12 hrs at 4 °C to cleave the His-tag. TEV-treated NMnPP was applied to a MonoQ ion-exchange column pre-equilibrated with 50 m*M* Tris-HCl, pH 8.0 and 5 m*M* β-mercaptoethanol. Then, linear gradient elution was performed up to 50 m*M* Tris-HCl, pH 8.0, 500 m*M* NaCl, and 5 m*M* β-mercaptoethanol. Pooled fractions were concentrated using Amicon centrifugal filters (Millipore) with a cut-off of 5 kDa and injected into HiLoad Superdex 200 prep grade chromatography (GE Healthcare) equilibrated with buffer C (20 m*M* Hepes-NaOH, pH 7.5, 150 m*M* KCl, 5 m*M* β-mercaptoethanol and 10% (*v*/*v*) glycerol). NMnPP protein fractions were examined by sodium dodecyl sulfate-polyacrylamide gel electrophoresis (SDS-PAGE, Fig. 3A), pooled, and concentrated at 15 mg ml^−1^ to do protein crystallization.

**Table 1.**
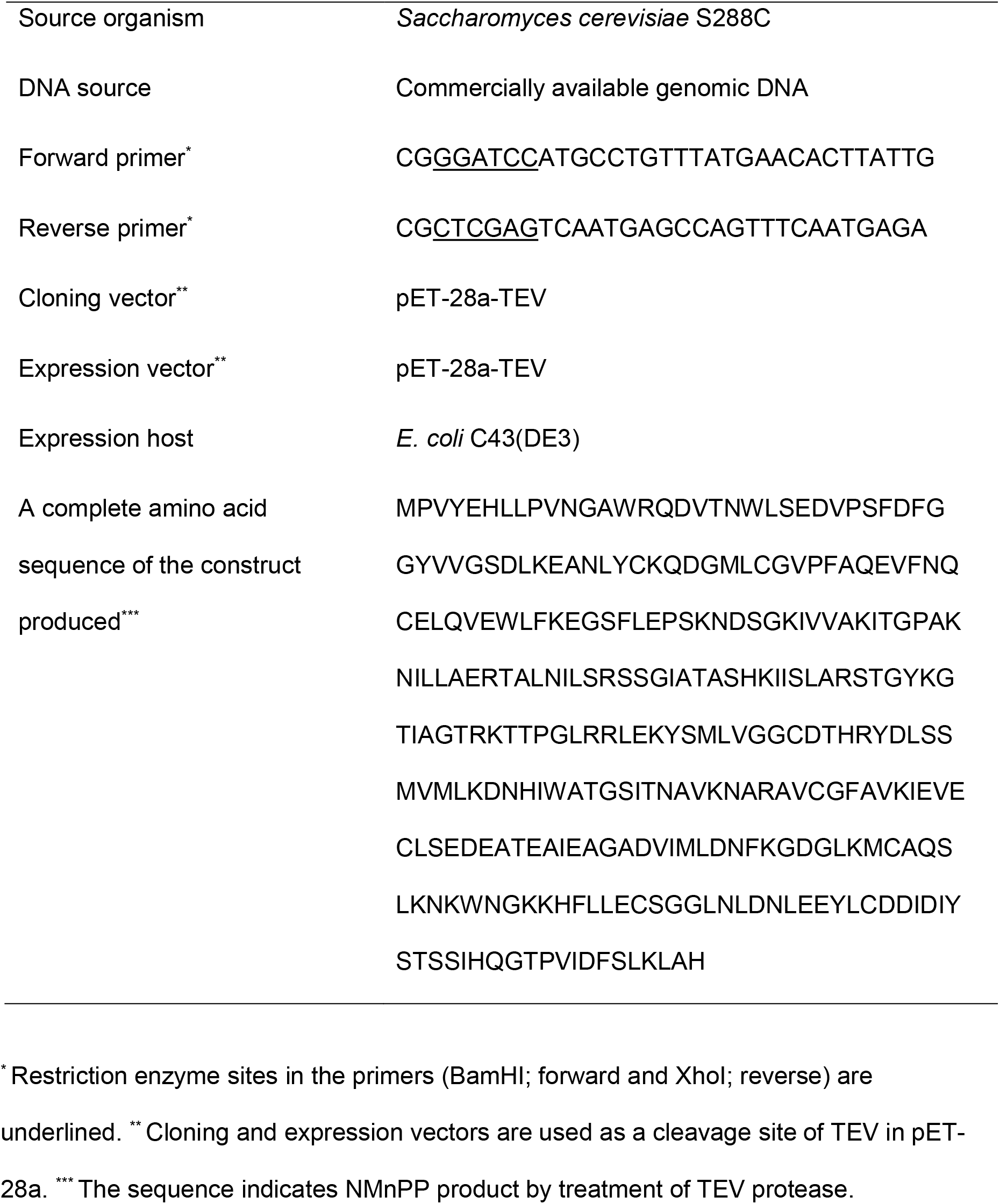
Macromolecule production information

**Table 2.**
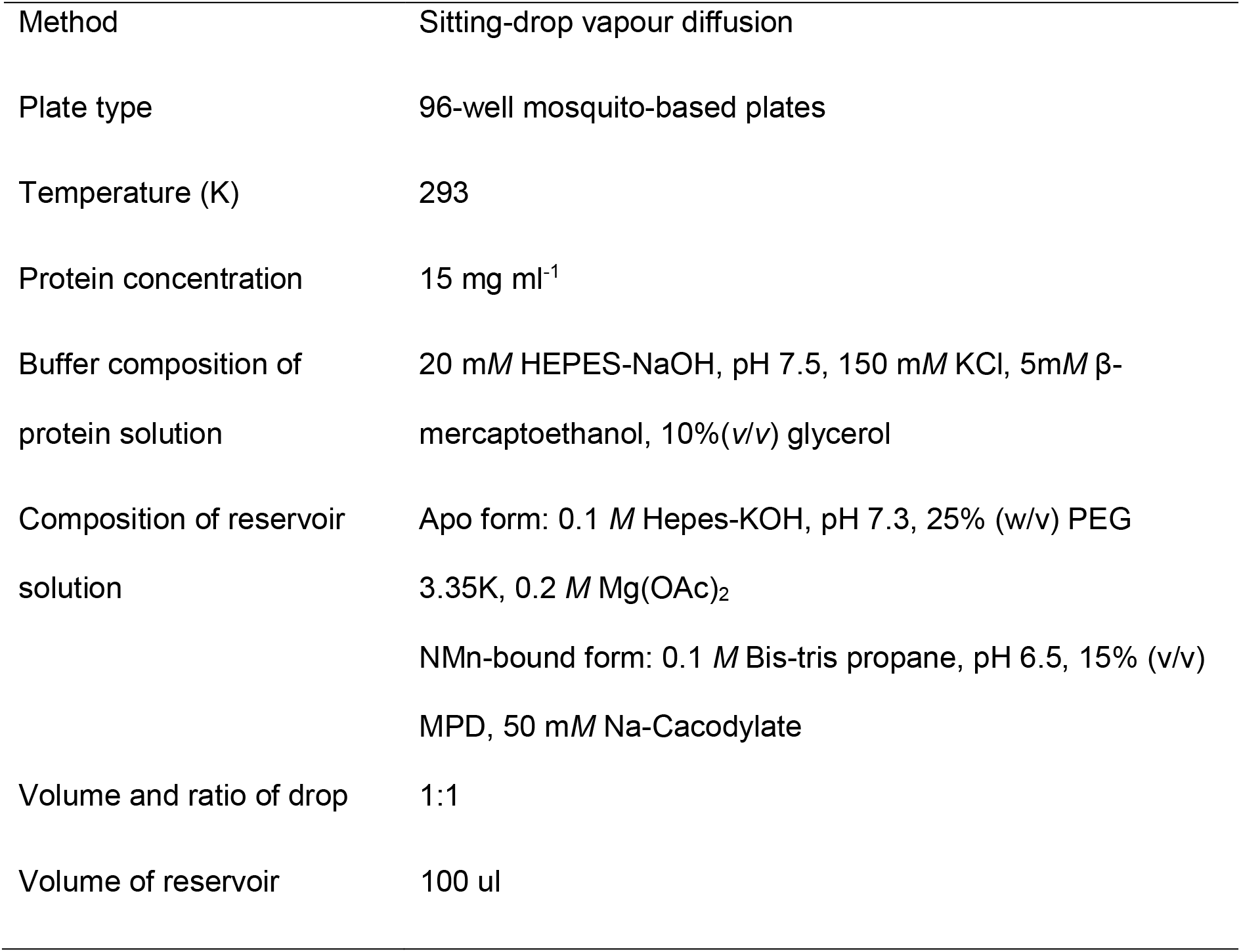
Protein crystallization

|  |  |
| --- | --- |
| Method | Sitting-drop vapour diffusion |
| Plate type | 96-well mosquito-based plates |
| Temperature (K) | 293 |
| Protein concentration | 15 mg ml<sup>-1</sup> |
| Buffer composition of protein solution | 20 mM HEPES-NaOH, pH 7.5, 150 mM KCl, 5mM β-mercaptoethanol, 10%(v/v) glycerol |
| Composition of reservoir solution | Apo form: 0.1 M Hepes-KOH, pH 7.3, 25% (w/v) PEG 3.35K, 0.2 M Mg(OAc)<sub>2</sub><br>NMn-bound form: 0.1 M Bis-tris propane, pH 6.5, 15% (v/v) MPD, 50 mM Na-Cacodylate |
| Volume and ratio of drop | 1:1 |
| Volume of reservoir | 100 ul |

### Crystallization

Apo NMnPP protein was initially crystallized using various screening kits from commercial company as Hampton Research, Creative Biostructure, and Jena Bioscience using an automatic crystallization machine Mosquito LCP (SPT Labtech). Based on the initial screening, both were further optimized as diffraction-qualified crystals using 96-well plates handled with the sitting-drop vapour diffusion method as same condition in Mosquito LCP machine applied with varying concentrations of buffer, precipitant, and metal ion. The NMn-bound NMnPP was incubated with an additional 0.15 *M* Zn(OAc)_2_ in buffer C and a 100-fold molar excess of R5P for 3 hrs on ice and then co-crystallized for two weeks at the same room temperature as the apo form. Apo NMnPP crystals were grown at 293 K in 0.15 μL hanging drops comprised of equal volumes of protein stock solution (~15 mg mL^−1^) and mother liquor containing 0.1 *M* Hepes-KOH, pH 7.3, 25% (w/v) PEG 3.35K, and 0.2 *M* Mg(OAc)_2_ (Fig. 3B). NMn-bound NMnPP crystals was grown in 0.15 μL drops comprised of equal volumes of protein stock solution (~15 mg mL^−1^) and mother liquor containing and 0.1 *M* Bis-tris propane, pH 6.5, 15% (v/v) MPD, and 50 m*M* Na-Cacodylate (Fig. 3C). Crystals grew to a maximum size of 0.15 × 0.05 × 0.05 mm (apo NMnPP) and 0.07 × 0.05 × 0.05 mm (NMn-bound form) over at least two weeks. Crystals were transferred into cryoprotectant (25% (*v*/*v*) glycerol in reservoir solution) and were frozen in liquid N_2_ at near 77K.

### Data collection and initial processing

Apo and NMnPP-bound crystals were harvested using cryo-loops and immersed promptly in cryo-protectant solution. X-ray diffraction data sets for apo and R5P-bound NMnPP crystals were collected at 1.74 and 1.99 Å resolution on beamline 5C at the Pohang Accelerator Laboratory, Republic of Korea. Exposure time was 1.0 s per frame. One complete data set was obtained with 1° of oscillation angle. X-ray diffraction data were collected at 100K. Data were indexed and scaled with HKL-2000 (Otwinowski *et al*., 1997). Crystallographic data statistics are presented in Table 3.

**Table 3.** Data collection and processing statistics

| Dataset | Apo | NCN-bound form |
| --- | --- | --- |
| X-ray beamline | PAL5C |  |
| Wavelength (Å) | 1.0000 |  |
| Rotation range per image (°) | 1 |  |
| Exposure time per image (s) | 1 |  |
| Space group | *R*32 | *R*32 |
| Unit cell dimensions<br>a,b,c (Å)<br>$\alpha,\beta,\gamma$ (°) | 155.313, 155.313, 67.507<br>90.000, 90.000, 120.000 | 155.091, 155.091, 69.204<br>90.000, 90.000, 120.000 |
| Resolution (Å) | 25.88-1.74(1.82-1.74) | 44.77-1.99(2.07-1.99) |
| Total reflections | 93582 | 75893 |
| Unique reflections | 1441(2484) | 20759(2451) |
| Completeness (%) | 99.3(96.4) | 95.9(91.4) |
| $R_{\text{merge}}^*$ (%) | 3.3(49.4) | 3.2(43.8) |
| Mean $I/\sigma(I)$ | 4.5(1.3) | 4.6(1.4) |
| CC1/2 (%) | 98.1(93.7) | 96.0(95.2) |

\* $R_{\text{merge}} = \sum_{hkl} \sum_i |I_i(hkl) - \langle I(hkl) \rangle| / \sum_{hkl} \sum_i I_i(hkl)$ , where $I_i(hkl)$ is the intensity of the $i$ th observation of reflection $hkl$ and $\langle I(hkl) \rangle$ is the average intensity of reflection $I_i(hkl)$ for all $i$ observation. Values in parentheses are for the highest resolution shell. The values in parentheses are the highest resolution in the shell.

### Negative-stain TEM analysis

For preparing apo and NMn-bound NMnPP on carbon EM grid (#U1013, EMJapan), we selected one carbon EM grid and place on the glass slide with the carbon side up (dark side of the grid up) using the fine tweezers and then placed the glass slide (containing the EM grid) in the PELCO easiGlow (Ted Pella). Treated EM grid by the self-closing tweezers were used to protein solution. Both proteins were diluted to a final concentration ranging from 0.015 to 0.05 mg/ml were placed onto freshly glow-discharged continuous carbon-coated side of carbon EM grids, stained with final 2% uranyl acetate for 20 sec, blotted by gently touching the edge of grid to filter paper, and then completely air-dried. Micrographs were recorded in a JEM-1230R transmission electron microscope operated at 100 kV and Gatan Ultracan digital camera, at a nominal magnification of 30,000.

## ACKNOWLEDGEMENTS

We appreciate the technical assistance of beamline 5C at Pohang Accelerator Laboratory for help with X-ray diffraction and data collection experiments. We gratefully acknowledge direct financial support from Gyeongbuk Institute for Bio industry and the Ministry of Trade, Industry, and Energy (P0009787).

## REFERENCES

Adams, P.D., Afonine, P.V., Bunkóczi, G., Chen, V.B., Davis, I.W., Echols, N., Headd, J.J., Hung, L.W., Kapral, G. J., Grosse-Kunstleve, R. W., McCoy, A. J., Moriarty, N. W., Oeffner, R., Read, R.J., Richardson, D.C., Richardson, J.S., Terwilliger, T.C., and Zwart, P.H. (2010). PHENIX: a comprehensive Python-based system for macromolecular structure solution. Acta Cryst D66, 213–221.

Amjad, S., Nisar, S., Bhat, A.A., Shah, A.R., Frenneaux, M.P., Fakhro, K., Haris, M., reddy, R., Patay, Z., Baur, J., Bagga, P. (2021). Role of NAD^+^ in regulating cellular metabolic signaling pathways. Mol Metabol 49, 101195.

Beal, M.F., Ferrante, R.J., Swartz, K.J., Kowall, N.W. (1991). Chronic quinolinic acid lesions in rats closed resemble Huntington’s disease. J Neurosci 11, 1649–1659.

Braidy, N., Guillemin, G.J., Mansour, H., Chan-Ling, T., Grant, R. (2011). Changes in kynurenine pathway metabolism in the brain, liver and kidney of aged female Wistar rats. FEBS J 278, 4425–4434.

Di Luccio, E., Wilson, D.K. (2008). Comprehensive X-ray structural studies of the quinolinate phosphoribosyl transferase (BNA6) from Saccharomyces cerevisiae. Biochemistry 47, 4039–4350.

Emsley, P., and Cowtan, K. (2004). Coot: model-building tools for molecular graphics. Acta Crystallogr D Biol Crystallogr 60, 2126–2132.

Guillemin, G.J., Brew, B.J., Noonan, C.E., Takikawa, O., Cullen, K.M. (2005a). Indoleamine 2,3 dioxygenase and quinolinic acid immunoreactivity in Alzheimer’s disease hippocampus. Neuropathol. Appl. Neurobiol. 31, 395–404

Guillemin, G.J., Wang, L., Brew, B.J. (2005b). Quinolinic acid selectively induces apoptosis of human astrocytes: potential role in AIDS dementia complex. J. Neuroinflammation 2, 16.

Kim, M.-K., Im, Y.J., Lee, J.H., Eom, S.H. (2006). Crystal structure of quinolinic acid phosphoribosyltransferase from Helicobacter pylori. Proteins 63, 252–255.

Kim, M.-K., Kang, K.B., Song, W.K., Eom, S.H. (2007). The role of Phe181 in the hexamerization of Helicobacter pylori quinolinate phosphoribosyltransferase. Protein J 26, 517–521.

Liu, H., Woznica, K., Catton, G., Crawford, A., Botting, N., Naismith, J.H. (2007). Structural and kinetic characterization of quinolinate phosphoribosyltransferase (hQPRTase) from homo sapiens. J Mol Biol 373, 755–763.

Matthews, B.W. (1968). Solvent content of protein crystals. J Mol Biol 33, 491–497.

McCoy, A.J. (2007). Solving structures of protein complexes by molecular replacement with Phaser. Acta Cryst D63, 32–41.

Opitz, C.A., Litzenburger, U.M., Sahm, F., Ott, M., Tritschler, I., Trump, S. (2011). An endogenous tumour-promoting ligand of the human aryl hydrocarbon receptor. Nature 478, 197–203.

Otwinowski, Z., and Minor, W. (1997). Processing of X-ray diffraction data collected in oscillation mode. Methods Enzymol 276, 307–326.

Sahm, F., Oezen, I., Opitz, C.A., Radlwimmer, B., von Deimling, A., Ahrendt, T., Adams, S., Bode, H.B., Guillemin, G.J., Wick, W, Platten, M. (2013). The Endogenous Tryptophan Metabolite and NAD+ Precursor Quinolinic Acid Confers Resistance of Gliomas to Oxidative Stress. Cancer Res 73, 3225–3234.

Vagin, A., and Teplyakov, A. (2010). Molecular replacement with MOLREP. Acta Cryst D66, 22–25.

Vega-Naredo, I., Poeggeler, B., Sierra-Sanchez, V., Caballero, B., Tomas-Zapico, C., Alvarez-Garcia, O., Tolivia, d., Rodriquez-Colunga, M.J., Coto-Montes, A. (2005). Melatonin neutralizes neurotoxicity induced by quinolinic acid in brain tissue culture. J Pineal Res 39, 266–275.

Winn, M.D., Ballard, C.C., Cowtan, K.D., Dodson, E.J., Emsley, P., Evans, P.R., Keegan, R.M., Krissinel, E.B., Leslie, A.G., McCoy, A., McNicholas, S.J., Murshudov, G.N., Pannu, N.S., Potterton, E.A., Powell, H.R., Read, R.J., Vagin, A., and Wilson, K.S. (2011). Overview of the CCP4 suite and current developments. Acta Cryst D67, 235–242.

Youn, H.-S., Kim, M.-K., Knag, G.B., Kim, T.G., Lee, J.-G., An, J.Y., Park., K.R., Lee, Y., Kang, J.Y., Song, H.-E., Park, I., Cho, C., Fukuoka, S., Eom, S.H. (2013). Crystal structure of Sus scrofa quinolinate phosphoribosyltransferase in complex with nicotinate mononucleotide. PLoS One 8, e62027.

Youn, H.-S., Kim, T.G., Kim, M.-K., Kang, G. B., Kang, J.Y., Lee, J.-G., An, J.Y., Park, K.R., Lee, Y., Lee, J.H., Eom, S.H. (2016). Structural Insights into the Quaternary Catalytic Mechanism of Hexameric Human Quinolinate Phosphoribosyltransferase, a Key Enzyme in de novo NAD Biosynthesis. Sci Rep 6, 19681.

